# KSHV uses viral IL6 to expand infected immunosuppressive macrophages

**DOI:** 10.1101/2023.03.05.531224

**Authors:** Michiko Shimoda, Tomoki Inagaki, Ryan R. Davis, Alexander Merleev, Clifford G. Tepper, Emanual Maverakis, Yoshihiro Izumiya

## Abstract

Kaposi’s sarcoma-associated herpesvirus (KSHV) is an oncogenic double-stranded DNA virus and the etiologic agent of Kaposi’s sarcoma and hyperinflammatory lymphoproliferative disorders. Understanding the mechanism by which KSHV increases the infected cell population is crucial for curing KSHV-associated diseases. Here we demonstrate that KSHV preferentially infects CD14^+^ monocytes and sustains viral replication through the viral interleukin-6 (vIL6)-mediated activation of STAT1 and 3. Using vIL6-sufficient and vIL6-deficient recombinant KSHV, we demonstrated that vIL6 plays a critical role in promoting the proliferation and differentiation of KSHV-infected monocytes into macrophages. The macrophages derived from vIL6-sufficient KSHV infection showed a distinct transcriptional profile of elevated IFN-pathway activation with immune suppression and were compromised in T-cell stimulation function compared to those from vIL6-deficient KSHV infection or uninfected control. These results highlight a clever strategy, in which KSHV utilizes vIL6 to secure its viral pool by expanding infected dysfunctional macrophages. This mechanism also facilitates KSHV to escape from host immune surveillance and to establish a lifelong infection. 160

**Summary:** KSHV causes multiple inflammatory diseases, however, the underlying mechanism is not clear. Shimoda et al. demonstrate that KSHV preferentially infects monocytes and utilizes virally encoded interleukin-6 to expand and deregulate infected monocytes. This helps the virus escape from host immune surveillance.

## Introduction

A virus is an infectious agent that can only replicate within a living host organism. Because of this dependence, viruses have evolved various mechanisms to exploit normal cell functions to escape from host immune surveillance for their survival advantage. This exploitation sometimes associates with prolonged damage to the host leading to pathologic processes and diseases caused by the viral infection (Hoenen and Groseth, 2022).

Kaposi’s sarcoma (KS)-associated herpesvirus (KSHV), or human gamma herpesvirus 8 (HHV-8), is an oncogenic double-stranded DNA virus that establishes a lifelong latent infection (Cesarman et al., 2019). KSHV is the etiologic agent of Kaposi’s sarcoma and is associated with two lymphoproliferative disorders: multicentric Castleman’s disease (MCD) and AIDS-associated primary effusion lymphoma (PEL). KSHV-inflammatory cytokine syndrome (KICS) may also represent a prodromic form of KSHV-MCD, which exhibits elevated KSHV viral loads and circulating inflammatory cytokines including IL-6, IL-10, and a KSHV-encoded IL-6 homolog (vIL-6) (Alomari and Totonchy, 2020; Caro-Vegas et al., 2020; Dumic et al., 2020; Goncalves et al., 2017; Polizzotto et al., 2012; Polizzotto et al., 2016). These highly inflammatory diseases are devastating, and a leading cause of cancer deaths in AIDS patients. Therefore, understanding the mechanism of KSHV infection is crucial for finding a cure for these diseases.

Natural transmission of KSHV most likely occurs through salivary and sexual transmission or during transplantation of KSHV-positive organs into a naïve recipient although initial KSHV infection is normally asymptomatic (Cesarman et al., 2019; Dittmer and Damania, 2019). In experimental settings, KSHV has been shown to infect various types of cell lines and primary cells such as epithelial cells and immune cells that include B cells, monocytes, and dendritic cells through binding to specific cell surface receptors such as Siglec DC-SIGN (Decker et al., 1996; Gregory et al., 2012; Rappocciolo et al., 2006; Staskus et al., 1997; van der Meulen et al., 2021). However, it remains unclear as to whether KSHV may preferentially infect a certain cell type among PBMC. The mechanisms by which KSHV facilitates a lifelong infection by increasing viral reservoirs and impacts the host immune system are also not entirely clear.

KSHV-encoded viral interleukin-6 (vIL6) is a homolog of human interleukin-6, which is encoded by KSHV ORF-K2 and is highly expressed during the lytic replication cycle (Sakakibara and Tosato, 2011). Viral IL6 is also expressed at physiologically functional levels in latently infected cells (Chandriani and Ganem, 2010) and is detectable in the sera and/or tumor tissues of patients with KS, PEL, and MCD (Aoki et al., 2001). Viral IL6 enhances cell proliferation, endothelial cell migration, and angiogenesis, leading to tumorigenesis, and has been suggested to be a driver of KICS (Giffin et al., 2015; Zhu et al., 2014). In addition, vIL6 transgenic mice develop IL6-dependent MCD-like disease (Suthaus et al., 2012) and supported tumor metastasis in a murine xenograft model (Fullwood et al., 2018). Mechanistically, vIL6 directly binds to the gp130 subunit of the IL6 receptor without the need of the IL6 receptor, α, and actives the JAK/STAT pathway to induce STAT3 phosphorylation and acetylation (Alomari and Totonchy, 2020; Cojohari et al., 2020; Molden et al., 1997). In addition, vIL6 activates the AKT pathway to promote numerous oncogenic phenotypes (Giffin et al., 2015; Morris et al., 2008; Morris et al., 2012; Zhu et al., 2014). STAT3 activation by vIL6 also increases the VEGF expression through the downregulating caveolin 1 (Alomari and Totonchy, 2020) and promotes angiogenesis, suggesting that vIL6 plays an important role in tumorigenesis through STAT activation. Given that the prototypical human IL6 plays a critical role in immune regulation and inflammation, vIL6 is thought to play a pivotal role in inflammatory KSHV diseases. Here we reveal the role of vIL6 in the regulation of monocytes by utilizing recombinant KSHV and de novo infection to the peripheral blood mononuclear cells.

## Results and Discussion

### KSHV preferentially infects CD14+ monocytes among PBMC and triggers an inflammatory response and macrophage differentiation

To study cell type-specific KSHV infection, we employed a single cell (sc)RNA-seq analysis approach. Recombinant KSHV (rKSHV.219) virions were purified by two serial ultracentrifugations from the culture supernatant of the iSLK.219 cell line, an inducible recombinant KSHV producer. Peripheral blood mononuclear cells (PBMCs) were infected with rKSHV.219 at MOI=1, fixed at various time points (day 0, 1, 2, and 4) after infection, and subjected to scRNA-seq analysis. KSHV infection and lytic replication in single cells were then monitored by the expression of K2 that encodes vIL6.

As shown in Fig. 1A, unsupervised, Uniform Manifold Approximation and Projection (UMAP) for dimension reduction analysis identified 9 clusters among peripheral blood mononuclear cells (PBMCs), each of which can be associated with known immune cell subsets based on corresponding lineage-specific gene expression. Thus, *KLRD1(CD94), CD14, APOBEC3A, VMO1, CD79A*, and *CD3E* genes were used as a marker for NK cells, monocytes, intermediate monocytes, non-classical monocytes, B cells, and T cells, respectively, based on the immune cell data available from the Human Protein Atlas (Uhlen et al., 2019). As shown in Fig. 1B, *de novo* KSHV infection of PBMCs in conjunction with scRNA-seq analysis at 1-day post-infection (dpi) revealed that K2 (vIL6) expression was almost completely overlapped with CD14 expression. It should be noted that scant K2 expression was also found in *CD79A*^+^ B cells, *KLRD1*^+^ NK and *CD3E*^+^ T cell clusters. KSHV has been shown to establish latent infection in B cells, causing B-cell lymphoma (Cesarman et al., 2019). However, the correlation in expression of CD14 and K2 was highly significant (p=8e-191) compared to that of CD19 and K2 (p=0.065) at 1 dpi. The following kinetic scRNA-seq analysis also showed that K2 and CD14 gene expression were colocalized during 1 to 4 dpi (Fig. 1C). In this experiment, the culture medium was not replenished during the period. Infectious KSHV virions that remained in the culture or were newly released from infected cells could have infected other cell subsets of PBMC besides CD14^+^ monocytes. Thus, KSHV selectively infects CD14^+^ monocytes and monocytes can support KSHV lytic replication along with vIL6 gene expression. The kinetic scRNA-seq analysis also revealed that KSHV infection in CD14^+^ monocytes triggered the activation and inflammatory response immediately after KSHV infection at 1dpi as evidenced by the upregulation of *IL3RA* (CD123), *CCL2*, and *TNFα* (Fig. 1C). The latter was followed by the expression of genes associated with differentiation of M2-like macrophages with a suppressive phenotype (*CD274/PD-L1*, *C1QA*, *RNASE1*, *IL-10*, and *CD163*) during 2-4 dpi (Fig. 1D). Collectively, scRNA-seq analysis demonstrates that KSHV preferentially infects CD14^+^ monocytes among PBMCs and triggers a monocyte inflammatory response followed by a macrophage differentiation program.

**Figure 1.**
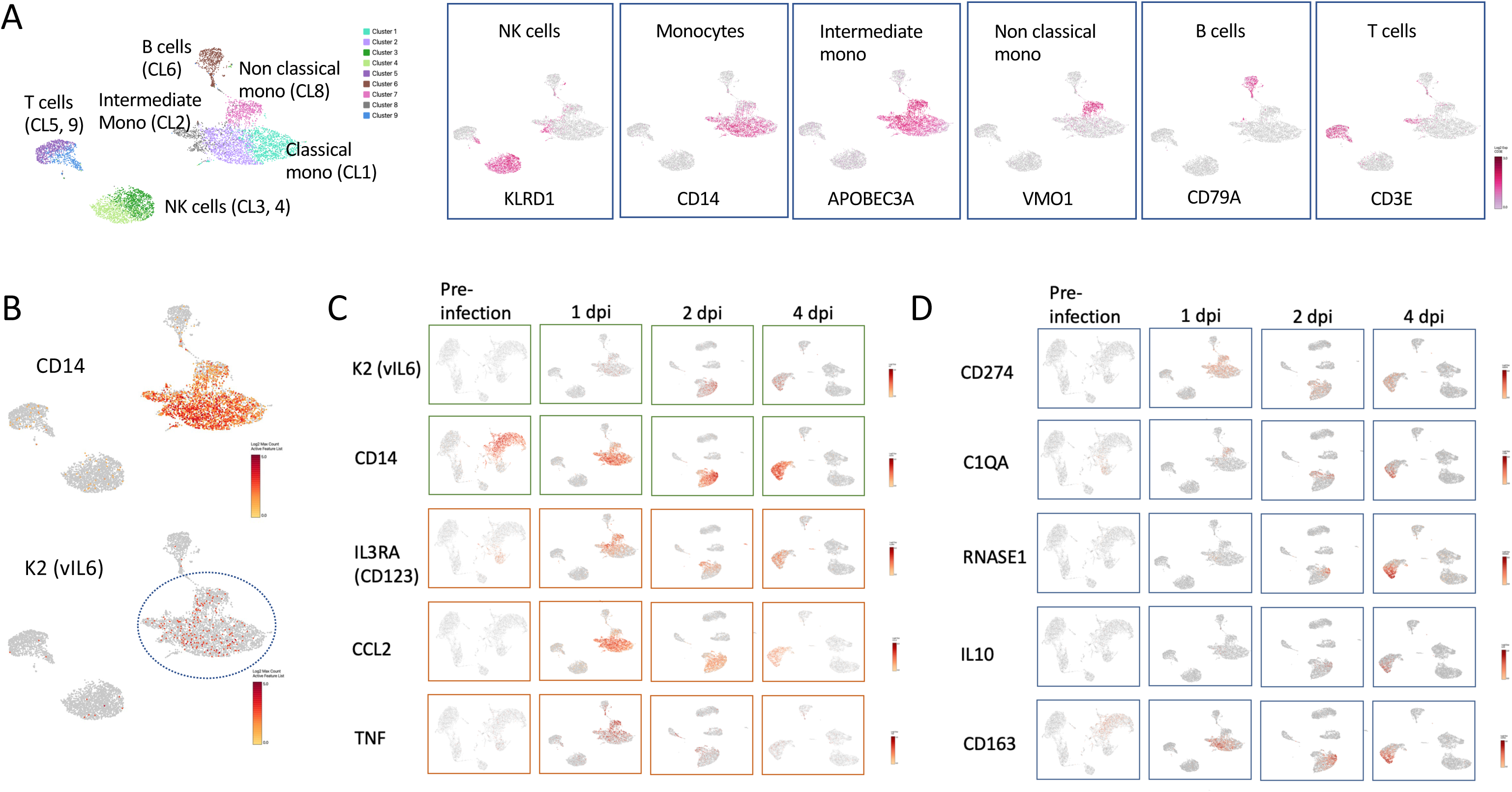
KSHV preferentially infects monocytes and triggers the monocyte inflammatory transcription program. PBMCs were infected with rKSHV.219 at MOI=1, fixed at various time points (pre-infection, day 1, 2, and 4) after infection, and subjected to 10x Genomics Chromium scRNA-seq analysis. (A) UMAP visualizes 9 PBMC clusters in a preinfectional sample based on lineage-specific gene expression of *KLRD1, CD14, APOBEC3A, VMO1, CD79A*, and *CD3E* genes for NK cells, monocytes, intermediate monocytes, non-classical monocytes, B cells, and T cells, respectively. (B) KSHV-encoded K2 (vIL6) expression overlapped with CD14 expression at 1 dpi. (C) The kinetic scRNA-seq analysis on pre-infection, 1 dpi, 2 dpi, and 4 dpi revealed that KSHV infection in CD14^+^monocytes triggered the activation and inflammatory response with the upregulation of *IL3RA* (CD123), *CCL2*, and *TNFα*. (D) Expression of genes in CD14^+^ monocytes associated with differentiation of M2-like macrophages with suppressive phenotype (*CD274/PD-L1*, *C1QA*, *RNASE1*, *IL-10*, and *CD163*). Gene expression levels are indicated by heatmap bars. A representative of three similar experiments is shown.

### KSHV infection of PBMC triggers STAT activation in dendritic cells and myeloid cell lineages

As a counterpart of the pleiotropic cytokine human IL6, viral IL6 has been shown to exhibit multifunctional pathologic roles in KS diseases (Sakakibara and Tosato, 2011). Therefore, we next studied the role of vIL6 in monocyte activation and differentiation using a *de novo* PBMC infection model. To this end, we generated recombinant KSHV lacking vIL6 expression by inserting stop codons in vIL6 (vIL6STOP) using the KSHV BAC clone BAC16. We also generated a revertant KSHV with intact vIL6 expression (vIL6REV) by changing the sequence back to the original wild-type sequence with BAC recombination. The lack of vIL6 expression in vIL6STOP was confirmed by Western blotting using a monoclonal antibody against vIL6 (a kind gift from Dr. Robert Yarchoan, NIH) (SFig. 1A).

First, to evaluate the signaling events triggered by KSHV infection in immune cells, PBMC from healthy donors (n=6, 3 males and 3 females) were infected with vIL6REV (with vIL6 expression) or vIL6STOP (without vIL6 expression) (MOI=1), or mock-infected (PBS). At 1 dpi, cells were fixed and subjected to CyTOF analysis using a 27-color signaling panel that can differentiate 25 immune cell subsets (SFig. 2). The expression levels of 10 signaling molecules were then determined in order to evaluate the early signaling events in PBMCs upon KSHV infection.

**Figure 2.**
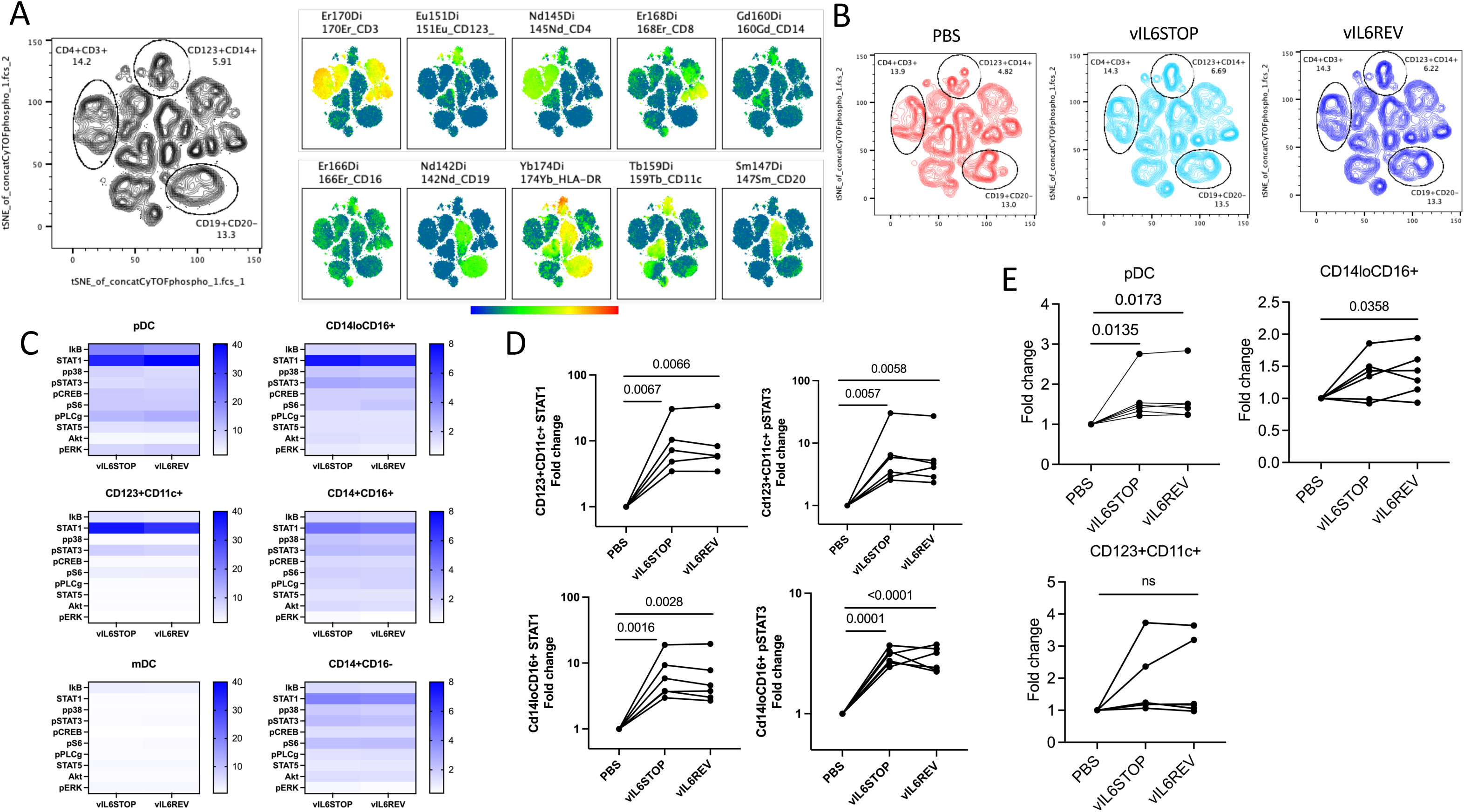
KSHV infection preferentially triggers STAT activation in monocytes and dendritic cells. PBMC from healthy donors (n=6) were infected with vIL6REV (KSHV with vIL6 expression) or vIL6STOP (KSHV without vIL6 expression) (MOI=1), or mock-infected (PBS). At 1 dpi, cells were fixed and subjected to CyTOF analysis. The expression levels of 10 signaling molecules were evaluated. Data combine two experiments (n=3 each). (A) tSNE for dimension reduction analysis was applied for each infection group after 1000 events from 6 samples in each group were concatenated. Immune cell subsets were clustered based on each lineage’s marker expression (Right panel). (B) The phenotypic changes induced in three immune cell populations corresponding to CD4^+^CD3^+^, CD19^+^CD20^-^, and CD123^+^CD14^+^ clusters by infection were identified by comparison between vIL6STOP, vIL-6REV, and the mock-infected (PBS) control group. (C) The activation status of 10 signaling molecules among plasmacytoid dendritic cells (pDC) and monocytic cell subsets including CD123^+^CD11c^+^ cells, myeloid (m)DC, CD14^lo^CD16^+^ non-classical monocytes, CD14^+^CD16^+^ intermediate monocytes, and CD14^-^CD16^+^ classical monocytes are shown in heat maps. Each subset was identified based on the lineage cell surface marker expression with gating strategies shown in SFig.2. (D) The fold change in the CyTOF signal intensity of STAT1 and pSTAT3 for CD123^+^CD11c^+^ cells (TOP panel) and CD14^+^CD16^+^ cells (Bottom panel) in infected groups compared to mock-infected (PBS) control group are shown. p values shown are by ratio paired t-test. p<0.05: statistically significant. (E) Individual percentages of pDCs, CD14^+^CD16^+^, and CD123^+^CD11c^+^ cells among DC gate for each PBMC sample (n=6) are compared between vIL6REV-and vIL6STOP-infected groups, and the uninfected group (PBS). p values shown are by ratio paired t-test. p<0.05: statistically significant.

Unsupervised t-distributed stochastic neighbor embedding (tSNE) for dimension reduction analysis was applied for each infection group. As shown in Fig. 2A, immune cell subsets were clustered based on each marker expression, and the phenotypic changes induced by infection were visualized and identified by comparison between vIL6STOP, vIL6REV, and the mock-infected (PBS) control group (Fig.2 B). The landscape of immune cell phenotypes visualized by the tSNE cell clustering was remarkably similar between the vIL6STOP and vIL6REV infection group, indicating that the anti-viral response was equally triggered in monocytes in vIL6STOP and vIL6-REV infection. Nonetheless, the tSNE visualization revealed that the phenotype of three immune cell populations corresponding to CD4^+^CD3^+^, CD19^+^CD20^-^, and CD123^+^CD14^+^ clusters as indicated in Fig. 2B was different between the infected groups and the mock-infected control group. The CD123^+^CD14^+^ cluster contains a heterogenous cellular population that also expresses CD11c, CD16, and HLA-DR (Fig. 2A). Given that the scRNA-seq results showed preferential KSHV infection in the CD14^+^ cell population in association with *CD123* (IL3R) upregulation (Fig. 1C), we considered that the changes observed in the CD123^+^CD14^+^ cluster represent the host cell anti-virus response in monocytes after KSHV infection. Changes in CD4^+^CD3^+^ and CD19^+^CD20^-^ clusters could be the result of infection and/or indirect activation such as through cytokine production from PBMCs in response to KSHV infection.

To evaluate signaling events in detail, we next used a knowledge-based analysis and identified 25 immune cell subsets based on the gating described in SFig.2. The activation status of each signaling molecule was evaluated among plasmacytoid dendritic cells (pDC) and monocytic cell subsets including CD123^+^CD11c^+^ cells, myeloid (m)DC, CD14^lo^CD16^+^ non-classical monocytes, CD14^+^CD16^+^ intermediate monocytes, and CD14^-^CD16^+^ classical monocytes (SFig.2), based on the fold change in the CyTOF signal intensity for infected groups compared to the mock-infected (PBS) control group (Fig. 2C and D). As expected, many of the 10 signaling molecules including STAT1, IKB, and phosphor-PLCψ were strongly upregulated in pDC (Fig. 2C), an immune cell subset responsible for the anti-viral response. Among CD11c^+^ DC subsets, STAT1 and phospho-STAT3 were the two most significantly activated pathways in CD123^+^CD11c^+^ cells, and less so in myeloid (m)DC (Fig. 2C and D). Among monocyte subsets, STAT1, phospho-STAT3, and pS6 were upregulated after infection (Fig. 2C and D). Consistent with the tSNE results, the activation profiles of 10 signaling molecules in pDC, DC, and monocytes were identical between vIL6STOP and vIL6REV infection (Fig. 2C and D). Thus, at 1dpi both vIL6STOP and vIL6REV viruses triggered indistinguishable host responses in pDC, DCs, and monocytes.

In association with upregulation of STAT 1 (Fig. 2C and D), the frequency of the pDCs among dendritic cells at 1dpi increased by infection regardless of the vIL6 expression (Fig. 2E) while the change in the frequency was not significant for CD123^+^CD11c^+^ cells. The frequency of the CD14^+^CD16^+^ population also increased by infection in the presence of vIL6 expression in KSHV (Fig. 2E). The vIL6-dependent response suggests a biological role of vIL6 in the activation of infected CD14^+^CD16^+^ monocytes.

### KSHV infection promotes activation and proliferation of CD14+ monocytes in a manner dependent on vIL-6

To further study the biological significance of vIL6 expression and STAT 1 and 3 activation, monocytes were isolated from PBMCs (n=6) using a magnetic beads-based negative enrichment method and infected with vIL6STOP or vIL6REV KSHV (MOI=1). The viability and phenotype were then analyzed by flow cytometry at 2 dpi. We found that total cell viability (Fig. 3A) and the frequency of activated CD274 (PDL-1) ^+^ cells (Fig. 3C) were significantly increased by KSHV infection regardless of the expression of vIL6. These results were expected based on the CyTOF signaling analysis results that demonstrate that the initial host response against vIL6REV and vIL6STOP infection was indistinguishable. As shown in Fig. 1, KSHV infection promptly induced inflammatory cytokine expression in monocytes. Inflammatory cytokines such as TNFα produced by activated monocytes can support their survival in an autocrine manner during the inflammatory response (Wolf et al., 2017). Therefore, it is possible that the effect of vIL6 on cell survival, if any, could be overridden by the effect of other inflammatory cytokines during the early host anti-viral response against KSHV.

**Figure 3.**
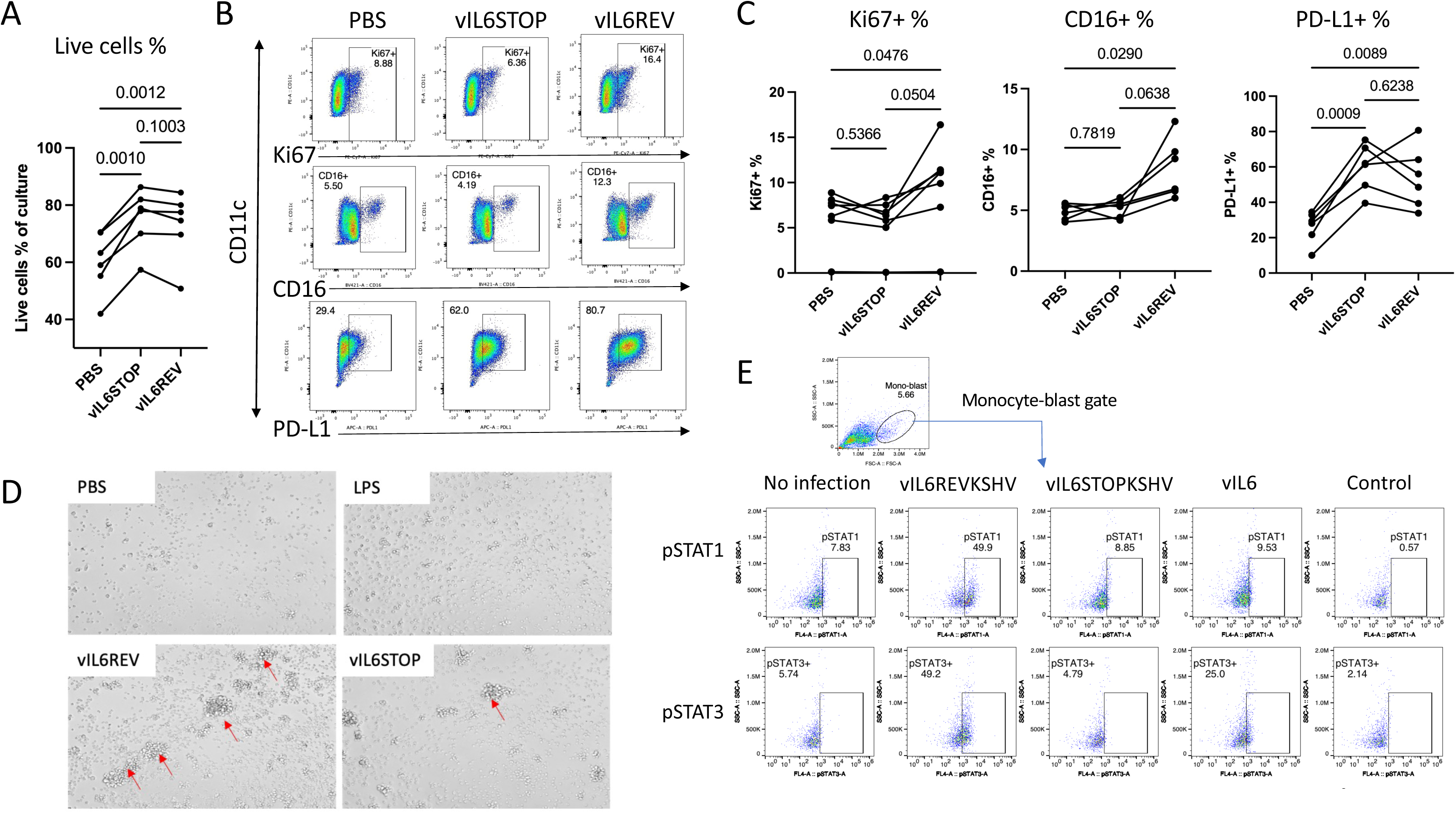
KSHV preferentially infects and expands CD14+ monocytes with an inflammatory response in a manner dependent on vIL6. Monocytes were isolated negatively from PBMCs (n=6) using magnetic beads-based negative enrichment and infected with vIL6STOP or vIL6REV rKSHV (MOI=1). (A) The frequencies of live cells at 2 dpi by live/dead staining are shown (B) Representative FACS profiles of infected monocytes analyzed by flow cytometry for Ki67, CD16, and PD-L1 expressing cells. (C) The frequencies for Ki67+, CD16+, and PDL-1+ cells are shown. Data represent two similar experiments. (D) Representative images of cell blasts in the indicated treatment conditions (x100). (E) Representative FACS profiles of each sample for pSTAT1 and pSTAT3 intracellular staining with the percentage of gated pSTAT1-and pSTAT3-expressing cells are shown. Data represent two similar experiments. *p* values shown are by ANOVA with a paired comparison with Tukey’s multiple comparison test. *p* < 0.05: statistically significant.

On the other hand, the frequency of Ki67^+^ proliferating cells and that of CD16^+^ cells were significantly increased in a manner dependent on the presence of vIL6 (Fig. 3B and C). In monocyte cultures, larger and more frequent proliferating blast cells were observed in response to vIL6REV infection compared to that of vIL6STOP infection (Fig. 3D). Intracellular staining of monocytes also confirmed that the frequency of pSTAT1 and pSTAT3 in monocytes was higher in vIL6REV-infection compared to that in vIL6STOP or mock-infection (Fig. 3E). These results collectively suggest that vIL6 expression during the early KSHV infection increases the proliferation of infected monocytes via STAT 1 and 3 activation. The cellular environment in monocytes may have a unique role in sustaining STAT activation with KSHV infection.

### KSHV infection changes the transcriptional landscape of macrophages in a manner dependent on vIL6 expression

To reveal the biological effect of vIL6 expression in infected monocytes, we next conducted a transcriptomic analysis of *de novo* infected monocytes. To this end, total RNA was isolated from monocytes harvested 7 days post KSHV infection and RNA-seq was performed (Fig. 4A in the Left panel). By 7 days post-infection, the recovery of vIL6REV-infected monocytes was higher than vIL6STOP-infected monocytes as shown later in Fig. 5A because of the increased Ki67^+^ proliferative responses (Fig. 3B and C). Principal component analysis (PCA) demonstrated markedly distinct transcriptional profiles between vIL6REV-infected and vIL6STOP-infected monocytes (Fig. 4A, the Right panel). Volcano plots depict the higher numbers of differentially regulated genes in vIL6REV-infected monocytes (Fig. 4B; e.g., relative to PBS control). To our surprise, vIL6STOP infection had little impact on the transcriptional landscape of monocytes by 7dpi, suggesting that vIL6 expression is needed for either sustaining cell signaling activation or maintaining KSHV lytic replication. Those two functions of vIL6 are likely to be mutually exclusive.

**Figure 4.**
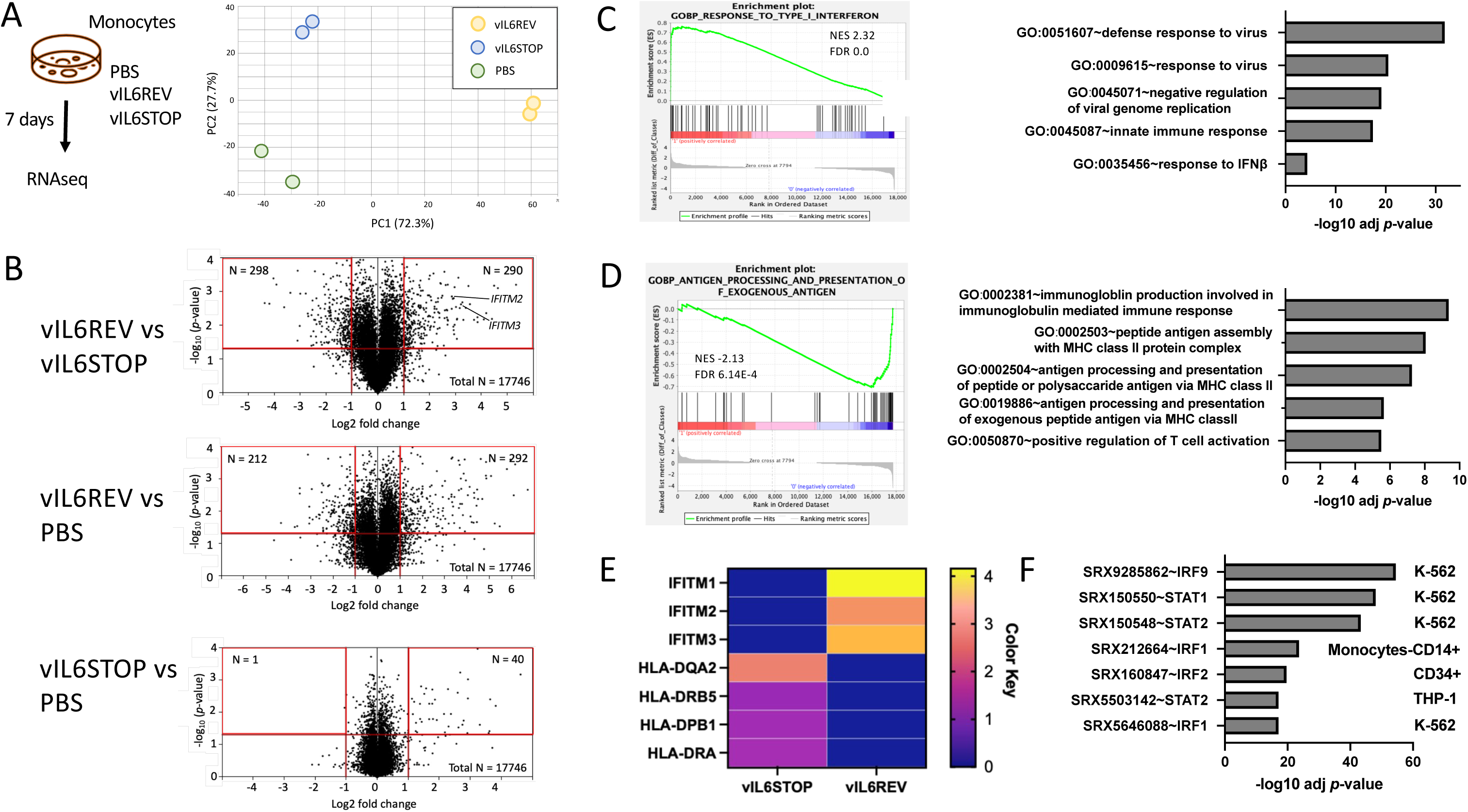
Differential gene expression analysis between wild-type KSHV and vIL6 deficient KSHV infected monocytes. (A) A schematic illustration of the KSHV infection experiment (Left). Monocytes were infected with vIL6REV, vIL6STOP, or mock-infected (PBS). At 7 dpi, cells were harvested for RNA-seq analysis (n=2/group). Principal component analysis (PCA) of RNA-seq data from independent biological replicates is shown (Right). The number in parenthesis indicates the percentage of total variance explained by each PC. Colored dots represent individual samples infected with mock (PBS-treated), vIL6REV, and vIL6STOP. (B) Volcano plots for differentially expressed genes (log2 FC threshold = 1, *p*-value threshold = 0.05) between vIL6REV and vIL6STOP (Top), vIL6 REV and mock (Middle) and vIL6STOP and mock (Bottom). The log2 FC indicates the mean expression for each. The functional enrichment analysis of upregulated genes (C) and downregulated genes (D) in vIL6REV-compared to vIL6STOP-infected monocytes. DAVID functional enrichment analysis was performed, and the analysis results of the top 5 enriched pathways are shown (Right). The representative results of Gene Set Enrichment Analysis (GSEA) are shown (Left). (E) Heatmap of representative genes from enriched gene sets for upregulated IFN-inducible genes (C) or downregulated MHC-II genes (D) in vIL6REV-compared to vIL6STOP-infected monocytes. (F) Putative transcription factors that are responsible for differentially expressed genes between vIL-6 REV and vIL-6 STOP KSHV infection. The bar charts indicate *p* values for enrichment. The following parameters were used; Organism: Homo sapiens (hg38), Experiment type: ChiP (TFs and others), Cell type Class: Blood, and Threshold for significance: 50.

**Figure 5.**
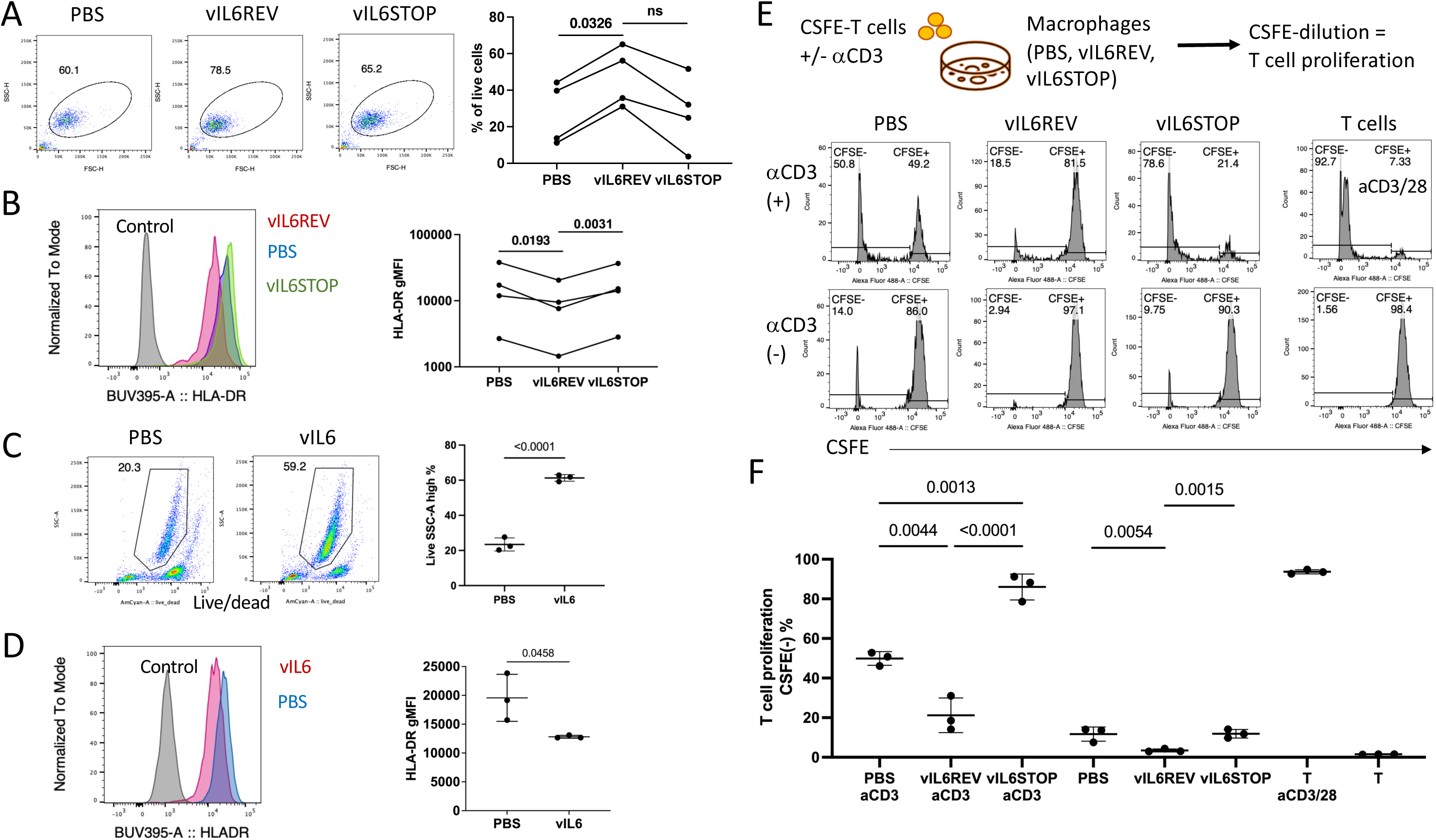
KSHV infection promotes macrophage differentiation with an immunosuppressive phenotype in a manner dependent on vIL6 expression. Monocytes were infected with vIL6REV or vIL6STOP (MOI=1), uninfected (PBS) and cultured for 7-14 days, or treated with vIL6 (200 ng/ml in serum-free medium) for 9 days. Each sample was analyzed by flow cytometry with isotype control staining as a control. (A) The average frequency of macrophages in each culture (n=3) is based on representative FSC-H and SSC-H profiles. Data combine four independent experiments with different donor samples. Dots represent the average % of biological replicates (n=2-3). (B) The average geometric mean fluorescent intensity (gMFI) of HLA-DR for each macrophage sample with representative overlays are shown. Data combine four independent experiments with different donor samples. Dots represent the average HLA-DR gMFI of biological replicates (n=2-3). (C) Representative FACS plots of live monocytes one week after vIL6 treatment and the average frequencies (n=3) are shown. (D) The average gMFI of HLA-DR for each macrophage sample in (C) with representative overlays are shown. Data represent two similar experiments. (E) An experimental design of T cell proliferation assay with macrophages (Top) and representative histograms of T cells (gated based on FSC-A and SSC-A) for CSFE are shown. T cells without or with anti-CD3/CD28 tetramer stimulation are used as controls. (F) The average frequencies of the CFSE-negative population determined as in (E) for each sample are shown (n=3). *p* values shown are by ANOVA with a paired comparison with Tukey’s multiple comparison test. *p* < 0.05: statistically significant. Data represent two similar experiments.

Among the genes differentially expressed between vIL6REV vs vIL6STOP infection, interferon-induced genes such as *IFITM2*, and *IFITM3*, are substantially upregulated in response to vIL6REV infection (Fig.4B, Top panel). Consistently, Gene Set Enrichment Analysis (GSEA) also revealed that genes associated with host anti-viral innate responses were enriched in vIL6REV infection (Fig. 4C, SFig. 3), while the pathways involving in antigen processing and presentation were downregulated in vIL6REV (Fig. 4D, SFig. 3). While we expected to see upregulation of IFN-related genes in KSHV infected cells, down-regulation of MHC class II genes are unexpected. Representative upregulated IFN-inducible genes and down-regulation of MHC-II were visualized in the heatmap (Fig. 4E). To identify the putative transcription factors that are responsible for vIL6-driven gene regulation, we examined transcription factors that are enriched on promoter regions of differentially transcribed genes. As shown in Fig. 4F, interferon regulatory factors, STAT1 and STAT2 were found as putative transcription factors that are responsible for driving vIL6-associated phenotypes.

### KSHV infection generates macrophages with an immunosuppressive phenotype in a manner dependent on vIL6 expression

Following the transcriptomic study that demonstrated transcriptional landscape changes in KSHV-infected macrophages, we next confirmed functional changes induced in monocytes after KSHV *de novo* infection on 7-14 dpi.

As shown above, in Fig. 1D, the M2-like macrophage differentiation program had been triggered by 4 dpi. Consistent with this, monocytes in cultures possessed features indicative of macrophages, such as being predominantly of large cellularity with higher FSC and SSC, as shown in Fig. 5A. Also, the recovery of cells was significantly increased in vIL6-REV infection compared to vIL6-STOP infection at 7-14 dpi. Consistent with the transcriptomic analysis in Fig. 4D, the macrophages derived from vIL6-REV infection (vIL6-REV Mac) expressed lower levels of HLA-DR with higher levels of CD16 compared to control groups (Fig. 5B). To test the direct effect of vIL6 in such phenotypic changes, we generated functional recombinant vIL6 capable of activating STAT1 (SFig. 1B and C). Monocytes were cultured in the presence of vIL6 (200 ng/ml in serum-free medium) for 9 days, and the phenotype was analyzed by flow cytometry. As shown in Fig. 5C, the overall monocyte viability was increased from 20% to 59% in the presence of vIL6. In addition, monocytes recovered from the 9 days culture showed a trend of reduced HLA-DR expression (Fig. 5D) compared to monocytes cultured without vIL6 (PBS). The results suggested that vIL6 treatment alone could induce phenotypic changes similar to vIL6REV infection.

Finally, to evaluate T cell co-stimulatory capacity, CSFE-labeled allogeneic T cells were cultured with macrophages derived from monocytes mock-infected or infected with vIL6REV or vIL6STOP. T cell proliferation was then evaluated as CSFE-dilution by flow cytometry (Fig. 5E, Top panel) in the presence or absence of anti-CD3 stimulation, and with the T cell culture with anti-CD3/CD28 tetramer stimulation as a positive control. As shown in Fig. 5E and summarized in Fig. 5F, the frequency of CFSE-negative proliferated T cells was significantly reduced in co-culture with macrophages from vIL6REV infection (18.5%) compared with uninfected (50.8%) or vIL6STOP-infection groups (78.6%) in the presence of anti-CD3 stimulation. Similar results were found in the cultures without anti-CD3 stimulation with the exception that the extent of T cell proliferation was smaller. These observations are consistent with the transcriptional analysis in Fig. 4, which collectively demonstrates that sustained vIL6-driven KSHV-lytic infection and continuous host anti-viral response in infected monocytes induces an immune suppressive phenotype in differentiating macrophages.

Monocytes play a key role in early infections by producing inflammatory cytokines and differentiating into subtypes that play distinct roles in the induction or resolution of inflammation (Watanabe et al., 2019)(Kapellos et al., 2019). Upon activation, some monocytes upregulate CD16 (FcgRIII), giving rise to CD14^+^CD16^+^ intermediate monocytes and CD14^-^CD16^+^ non-classical monocytes, with a prolonged circulating lifespan. These CD16^+^ monocytes normally exhibit high HLA-DR expression for the capability of antigen presentation, transendothelial migration, and FcR-mediated phagocytosis for anti-viral response (Yeap et al., 2016). CD16^+^ monocytes also express the highest levels of TNFR2 (CD120b) (Hijdra et al., 2012), which can upregulate the anti-inflammatory cytokine IL-10 cytokine (Gane et al., 2016) and supports either cell survival or death (Webster and Vucic, 2020). In this context, we observed an increase in CD14^+^CD16^+^ cells during early *de novo* KSHV infection (Fig.2E and 3B). We speculate that KSHV infects CD14^+^CD16^+^ monocytes by exploiting their FcR-and complement-mediated phagocytosis mechanism and expands them. The vIL6-driven sustained STAT1/3 activation may provide infected monocytes with a proliferative advantage over uninfected cells to increase their population size. Furthermore, vIL6-driven lytic infection constitutively triggers host anti-viral responses such as IFN-pathway activation, which in turn, induces regulatory mechanisms (McNab et al., 2015). In the case of *Mycobacterium*-infected macrophages, Type-I IFN signaling decreases energy metabolism (i.e. glycolysis and mitochondrial functions) (Olson et al., 2021) and causes cell death (Zhang et al., 2021). In *Listeria-*infected macrophages, stress-induced p38 mitogen-activated protein kinases (MAPK) signaling synergistically enhances IFN-stimulated genes (ISGs). In macrophages, IL-10 production as a part of the IFN-induced regulatory mechanism has been shown to dampen the capability of T cell activation and antigen presentation (McNab et al., 2014; Redpath et al., 1999; Zdrenghea et al., 2015). We show that the induction of a suppressive M2-like macrophage gene program was induced as early as 4 dpi in our scRNA analysis (Fig. 1). Consistent with this, earlier studies by other researchers also showed that KSHV infection and STAT3 activation induce a suppressive phenotype in dendritic cells and the THP-1 cell line (Bhaskaran et al., 2017; Campbell et al., 2014; Gilardini Montani et al., 2020; Qin et al., 2010). Accumulation of such immunosuppressive macrophages may increase the risk of secondary infection or cause chronic infection and inflammation in the host by dysregulating immune responses; this mechanism may increase the risk of KSHV-associated disease development.

In summary, this study demonstrates the preferential KSHV infection of monocytes and highlights the critical role of vIL6 in expanding infected monocytes that facilitate a long-term virus reservoir. Similar hijacking strategies to exploit monocyte development and functions have been also reported in Cytomegalovirus (Baasch et al., 2021; Baasch et al., 2020; Cojohari et al., 2020; Shnayder et al., 2020). The current study was unique in that vIL6STOP and vIL6REV rKSHV infection system was introduced to delineate the role of vIL6 from that of the other KSHV-encoded multifunctional factors; however, this study does not clarify whether vIL6 is directly involved as a driver in the expression of ISGs under the host defense program in infected monocytes. Nonetheless, vIL6 was required for the sustaining KSHV lytic infection in monocytes. As shown in Fig.4, in the absence of vIL6, KSHV infection in monocytes generated macrophages indistinguishable with those from uninfected monocytes. In this context, our study suggests that vIL6 and monocytes/macrophages are important therapeutic targets in KSHV-associated diseases. Understanding the mechanism in which vIL6 overcomes the IFN-induced regulatory mechanism and promotes cell proliferation would help develop therapies for KSHV infection. It is also important to determine whether KSHV establishes latent infection in macrophages to facilitate a life-long viral reservoir in patient samples. Of note, KSHV is found in circulating monocytes and macrophages (Decker et al., 1996; Rappocciolo et al., 2006), and blood-derived KS-like spindle cells were found to express macrophage markers (Uccini et al., 1997). Future studies are needed to investigate the possible role of monocytes/macrophages as a long-term viral reservoir in patients with KSHV-associated diseases. (16894w/o space)

## Materials and Methods

### Cells

LRCs from healthy donors were purchased from Vitalant. PBMCs were prepared by a standard Ficoll gradient method. PBMCs with a 10-30% range of CD14^+^ monocytes content measured by flow cytometry were stored in liquid N2 and used for experiments. PBMCs were thawed in warm media, washed twice, counted with the Countess automated cell counter (ThermoFisher), and resuspended at 2-5×10^6^ viable cells/ml. Monocytes were isolated with EasySep^TM^ Human monocyte isolation kit (STEMCELL) according to the manufacturer’s protocol. PBMCs and CD14^+^ monocytes were cultured in RPMI1640 medium containing 10% FBS plus antibiotics (2-5 x 10^6^ cells/ml, 200uL/well in a 96 well plate, 400uL/well in a 48 well plate, or 1ml/well in a 24 well plate) supplemented with or without stimuli for various time as described in each figure legend.

### Virus preparation

Doxycycline-inducible rKSHV.219 producer iSLK.219 cell lines were maintained in DMEM supplemented with 10% FBS, 1% penicillin-streptomycin-L-glutamine solution, 10 μg/mL puromycin, 400 μg/mL hygromycin B, and 250 μg/mL G418. For virus preparation, iSLK.219 cell lines were stimulated with 0.3 mM Sodium Butylate and 1 μg/ml Doxycycline for 5 days. Recombinant KSHV particles were then purified after two serial ultracentrifugation steps from the culture supernatant of activated iSLK.219 cell lines. Briefly, the culture supernatant was centrifuged at 300 × g for 10 min, and then passed through a 0.8-μm filter to remove cellular debris, and then viral particles were concentrated by ultracentrifugation at 25,000 rpm for 2 hrs at 4°C with a Beckman SW28 rotor. The viral precipitates were resuspended in 500 μL of DMEM along with the residual ∼500 μL media in the centrifuge tube and stored at −80°C until use.

### Quantification of viral copy number

Two hundred microliters of cell culture supernatant were treated with DNase I (12 μg/mL) for 15 min at room temperature to degrade unencapsidated DNA. This reaction was stopped by the addition of EDTA to 5 mM followed by heating at 70°C for 15 min. Viral genomic DNA was purified using the QIAamp DNA Mini Kit according to the manufacturer’s protocol and eluted in 100 μL of buffer AE. Four microliters of the eluate were used for real-time qPCR to determine viral copy number, as described previously (Izumiya et al., 2009).

### Recombinant KSHV

To test the role of vIL6, rKSHV.219 lacking vIL6 protein expression (vIL6STOP) was generated by adding stop codons in the vIL6 reading frame with BAC16 (vIL6STOP). We also generated a revertant rKSHV.219 with vIL6 protein expression (vIL6REV) by changing the sequence of vIL6STOP back to the original wildtype sequence with BAC-recombination.

### KSHV *de novo* infection

KSHV infection was conducted at MOI=1. Briefly, PBMCs or monocytes (1-2 x 10^6^ cells) were suspended in RPMI 1640 medium containing 10% intact FBS (not heat inactivated) supplemented with 8 ug/ml polybrene, and an appropriate amount of rKSHV.219 in PBS (less than 10% of total volume) or PBS alone (i.e., as mock infection) was added to the culture. Cells were kept in the same culture medium until analysis.

### Single-cell RNA-seq

2 x 10^6^ PBMCs were infected with rKSHV.219 at MOI=1, washed and fixed with paraformaldehyde according to the 10x genomics cell preparation guide (CG000053) at various time points (day 0, 1, 2, and 4) after infection. Single-cell suspensions (10^4^ cells) were submitted to the UC Davis Comprehensive Cancer Center (UCDCCC) Genomics Shared Resource for processing on the Chromium Controller (10x Genomics) for cell partitioning into individual GEMs (gel bead-in-emulsion), generation of barcoded cDNAs with unique molecular identifiers (UMIs), and construction of barcoded sequencing libraries were constructed using the Chromium Single Cell 3’ Reagent Kits v2 (10x Genomics). The libraries were then multiplexed and sequenced (∼250 million mapped reads per sample) on an Illumina NovaSeq 6000 sequencing system. Single-cell data were analyzed with the Cell Ranger v2.1 pipeline (10x Genomics). The pipeline included alignment to the hg38 human reference genome and human herpesvirus 8 strain (GQ994935.1) reference genome with STAR (Dobin et al., 2013), Uniform Manifold Approximation and Projection (UMAP), and K-means clustering. Read count matrices obtained from the pipeline were normalized using the log normalization method from the “Seurat” R package (Stuart et al., 2019). We investigated the association between genes using the Pearson product-moment correlation coefficient. To compute the correlation coefficient and p-value, we utilized the cor.test() function from the stats package in R (R. Core Team, 2021).

### mRNA-seq

RNA was prepared from infected monocytes using a QIAGEN RNeasy Miniprep kit according to the manufacturer’s protocol and submitted to the UCDCCC GSR for mRNA-seq analysis. Indexed, stranded mRNA-seq libraries were prepared from total RNA (100 ng) using the KAPA Stranded mRNA-Seq kit (Roche) according to the manufacturer’s standard protocol. Libraries were pooled and multiplex sequenced on an Illumina NovaSeq 6000 System (150-bp, paired-end, >25 × 10^6^ reads per sample). The RNA-Seq data was analyzed using a STAR-HTSeq-DESeq2 pipeline. Raw sequence reads (FASTQ format) were mapped with STAR to the reference human genome assembly (GRCh38/hg38, GENCODE release 36) and quantified with HTSeq (Patro et al., 2017). The resulting data were first filtered by average normalized count data >1 and volcano plots production and listing differentially expressed genes was performed by Subio Platform ver. 1.24 (Subio Inc., Amami, Japan). For transcription factor analysis, differentially expressed genes were submitted to Chip-Atlas to analyze common regulators and to predict transcription factor binding (Oki et al., 2018).

### Phosphoflow PBMC CyTOF

This assay was performed by the Human Immune Monitoring Center at Stanford University. 2 x 10^6^ PBMCs were infected with rKSHV.219, washed twice with PBS and then fixed with paraformaldehyde. Cells were washed twice with CyFACS buffer (PBS supplemented with 2% BSA, 2 mM EDTA, and 0.1% sodium azide) and stained for 30 min at room temperature with 20 mL of surface antibody cocktail. Cells were washed twice with CyFACS, permeabilized with 100% methanol, and kept at −80C overnight. The next day, cells were washed with CyFACS buffer and resuspended in a 20 mL intracellular antibody cocktail in CyFACS for 30 min at room temperature before washing twice in CyFACS. Cells were resuspended in a 100 mL iridium-containing DNA intercalator (1:2000 dilution in 2% PFA in PBS) and incubated at room temperature for 20 min. Cells were washed once with CyFACS buffer and twice with MilliQ water. Cells were diluted to 750×10^5^ cells/mL in MilliQ water and acquired on CyTOF. Data analysis was performed using FlowJo v10.8.1 by gating on intact cells based on the iridium isotopes from the intercalator, then on singlets by Ir191 vs cell length followed by cell subset-specific gating as described in SFig. 2.

### Flow cytometry

mAbs and isotype-matched controls were purchased from BD Bioscience (BUV395 conjugated HLA-DR, BUV395 conjugated to CD11c, PerCP-Cy5.5 conjugated to CD123, PE conjugated to CD11c, Alexa Fluor 647 conjugated to pSTAT-1, and Alexa Fluor 647 conjugated to pSTAT-3), and Biolegend (APC conjugated CD163, BV421 conjugated to CD16, PE-Cy7 conjugated to Ki67, APC conjugated to PD-L1, APC conjugated to CD14, and Alexa Fluor 488 conjugated to CD16). Cells in culture were washed twice with PBS and resuspended in FACS buffer (PBS supplemented with 1% FBS) at 1×10^7^ cells/ml. 50 uL cells per well were stained with Aqua LIVE/DEAD cell viability dye (Invitrogen, Carlsbad, CA) according to the manufacturer’s instructions, washed with FACS Buffer, and then stained for 45 min at room temperature with antibodies. For intracellular staining of pSTAT1 and pSTAT3, cells were fixed with paraformaldehyde, permeabilized with methanol, and kept at −80 C overnight. The cells were washed with FACS buffer and stained with pSTAT-1 Alexa Fluor 647 and pSTAT-3 Alexa Fluor 647. Cells were washed three times with FACS buffer and resuspended in 200 uL FACS buffer. For typical experiments, 10^5^ PBMCs per sample were collected using DIVA 6.0 software on a Fortessa flow cytometer (BD Biosciences) at the UCDCCC Flow Cytometry Shared Resource. Data analysis was performed using FlowJo v10.8.1 by gating on live cells based on forward versus side scatter profiles, then on singlets using forward scatter area versus height, followed by dead cell exclusion using Aqua LIVE/DEAD viability dye, and then cell subset-specific gating.

### T cell proliferation assay

Monocytes were infected with vIL6REV or vIL6STOP, or mock-infected (PBS). Macrophages derived from KSHV-infected monocyte cultures were recovered at 7 dpi after being washed with PBS twice followed by gentle pipetting and suspended in RPMI1640 medium containing 10% FBS. T cells were isolated from PBMCs using the EasySep^TM^ Human CD3 positive selection kit II (STEMCELL) according to the manufacturer’s protocol. T cells were stained in PBS with the CellTrace^TM^ CFSE Cell Proliferation Kit (ThermoFisher) according to the manufacturer’s protocol and suspended in RPMI1640 medium containing 10% FBS. CSFE-labeled CD3+ T cells (10^4^) were incubated with 10^3^ macrophages from KSHV-infected monocyte cultures in 200 uL/well in a 96-well U-bottom plate, without or with 100 ng/ml anti-CD3 antibody (BD Bioscience) for 5 days. For positive and negative controls, T cells were stimulated with ImmunoCult Human CD3/28 T cell activator (Stemcell) or left unstimulated. Cells were recovered from cultures and analyzed using a BD Accuri^TM^ flow cytometer. Data analysis was performed using FlowJo v10.8.1 by gating lymphocytes based on forward versus side scatter profiles. The average frequencies of the CFSE-negative population were determined based on the negative control.

### Statistical analysis

Statistical analyses were performed using GraphPad Prism 9.4.1 software. Results are shown as mean ± SD with dots represent individual measurements. Statistical significance was determined by Student’s t-test, ratio paired t-test, or one-way ANOVA with Tukey’s multiple comparison test, and correction for false discovery rate (FDR) as described in each Figure legend. FDR corrected p < 0.05 was considered statistically significant.

### Online supplemental material

SFig.1 shows the characterization and validation of recombinant vIL6STOP and vIL6REV KSHV and recombinant vIL6 used in this study. SFig.1A confirms the presence or absence of vIL6 expression in vIL6REV or vIL6STOP iSLK.219 cell lines that were used for virus preparation, respectively, by Western blotting. SFig.1B shows the identity and purity of recombinant vIL6 and its functionality to activate STAT1/3 in 293T and iSLK cell lines by Western blotting using a loss-of-function vIL6 mutant as a control. SFig.2 illustrates the CyTOF phospho-panel markers and the gating strategy used for analysis of immune cell subsets. SFig.3 shows GSEA analysis of RNAseq data comparing vIL6REV-infected and uninfected macrophages with charts displaying the top 5 enriched upregulated and down regulated pathways.

### Data availability

The scRNA-seq and RNA-seq data underlying Figs. 1 and 4 are openly available at GEO NCBI under accession number GSE227167. The data underlying Fig.2 are available in 10.6084/m9.figshare.22273279). The data underlying Figs. 3 and 5 are available in the published article and its online supplemental material. The reagents and cell lines are available upon request from Y. Izumiya.

## Acknowledgment

The authors would like to thank Dr. Robert Yarchoan (National Cancer Institute, USA) and Dr. Harutaka Katano (National Institute of Infectious Disease, Japan) for the generous gift of rabbit antibodies against vIL6. We would like to thank Andreas Koni for his assistance in preparing the manuscript. We would like to thank Immune Modeling, Analysis and Diagnostic, Genomics, and Flow Cytometry Shared Resources of the UC Davis Comprehensive Cancer Center Shared Resources for expert support with 10x Genomics scRNA-seq, total RNA-seq, and flow cytometry analyses, and the Stanford Human Immune Monitoring Center for expert CyTOF analysis. M.S. and Y.I. are founders of VGN BIO, Inc. The authors have no additional financial interests.

This work was supported by grants from the National Cancer Institute CA232845-02S1 (M. Shimoda), CA225266, CA232845, and AI167663 (Y. Izumiya), and UC Davis Dermatology Seed grant (M. Shimoda). M. Shimoda was supported by the National Cancer Institute/CRCHD/CURE program. The Immune Modeling, Analysis and Diagnostic, Genomics, and Flow Cytometry Shared Resources are supported by the UC Davis Comprehensive Cancer Center Support Grant (CCSG) awarded by the National Cancer Institute (NCI P30CA093373).

## Author contributions

Conceptualization: M. Shimoda, and Y. Izumiya. Experimental design: M. Shimoda and Y. Izumiya. Experiment and Data acquisition and analysis: M. Shimoda, R. Davis, A. Merleev, T. Inagaki, and Y. Izumiya. Funding acquisition: M. Shimoda and Y. Izumiya. Manuscript preparation: M. Shimoda. Review & Editing: M. Shimoda, R. Davis, A. Merleev, C.G. Tepper, E. Maverakis, T. Inagaki, and Y. Izumiya.

---

**SFig1.**
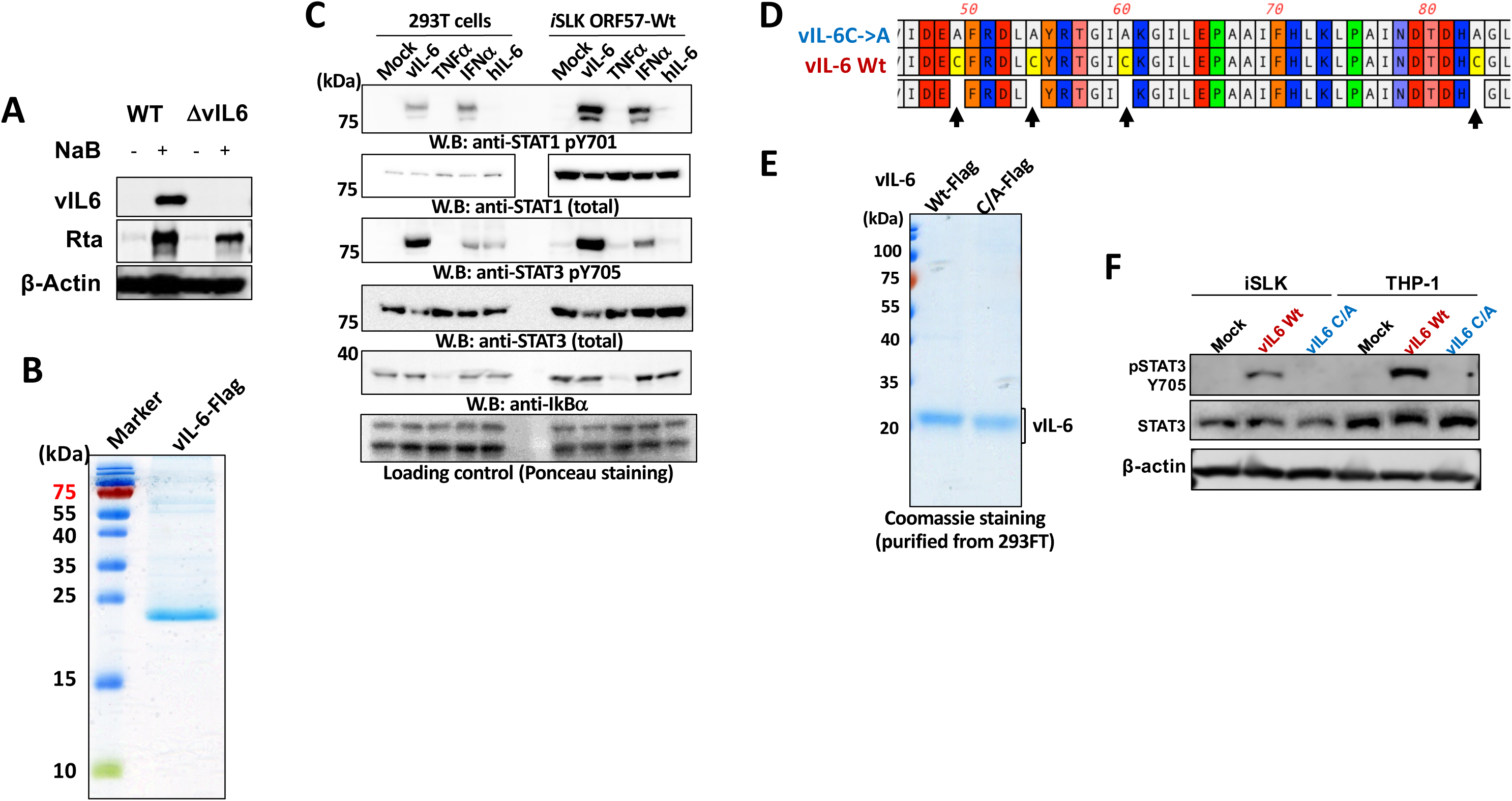
Validation of recombinant (r)KSHV and vIL6. (A) Lack of vIL6 protein expression in vIL6 STOP KSHV producing iSLK cells. iSLK cell lines transduced to express vIL6 of wild type (WT: vIL6 REV) or deficient (1¢1vIL6: vIL6 STOP) rKSHV were stimulated with Sodium Butyrate to induce KSHV lytic activation with vIL6 and Rta expression. KSHV-encoded vIL6 and Rta protein, and host β-actin as a control was detected by Western blotting with specific antibodies. (B) Recombinant vIL6-Flag protein was produced in insect cell lines and the purity confirmed by Western blotting with rabbit anti-vIL6 monoclonal antibody. A gel image with molecular weight markers is shown. (C) 293T and iSLK ORF57-wt cell lines were stimulated with 200ng/ml recombinant vIL6 along with controls, TNFα, IFNα, and human (h)IL6, or mock treated. STAT1 and 3 activation was studied by Western blotting with specific antibodies for total STAT1, STAT1 pY701, total STAT3, STAT3 pY705, and IkBα as controls. Ponceau staining for each well are shown as loading control. (D) To validate vIL6 activity, wild type (Wt) or mutant (C->A) are generated. Arrows indicate the location of amino acid residues where mutations are introduced. (E) Wt and C/A mutant vIL6 proteins were expressed in 293FT cell lines, the purity of which confirmed by Western blotting. (F) iSLK and THP-1 cell lines were stimulated with 200ng/ml recombinant Wt or C/A mutant vIL6, or mock treated. STAT3 activation was studied by Western blotting with specific antibodies for total STAT3, STAT3 pY705, and β-actin as controls. Wt but not C/A mutant was able to trigger STAT3 activation.

**SFig2.**
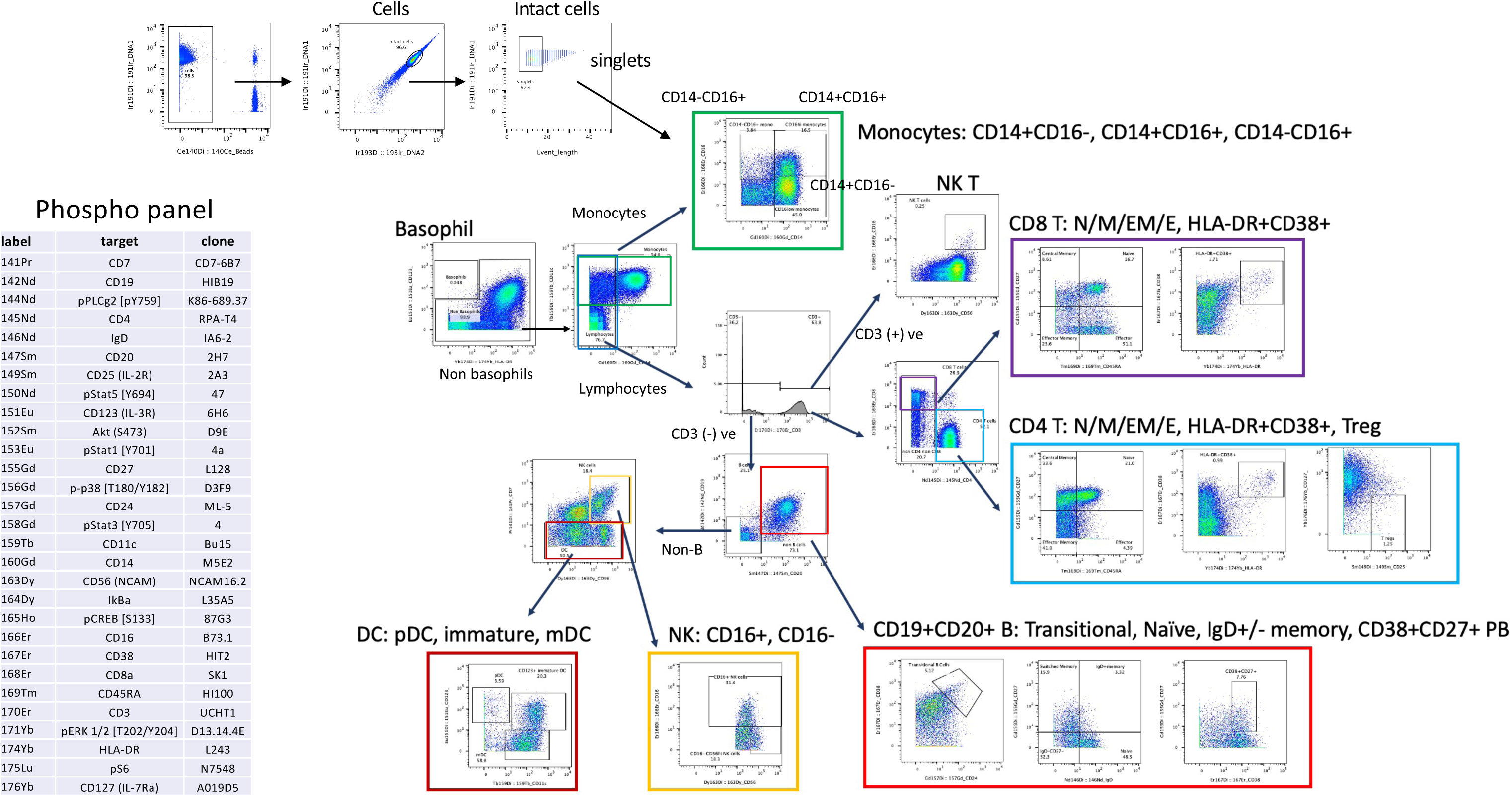
CyTOF phosphor panel and the gating strategy to identify immune cell subsets are shown.

**SFig3.**
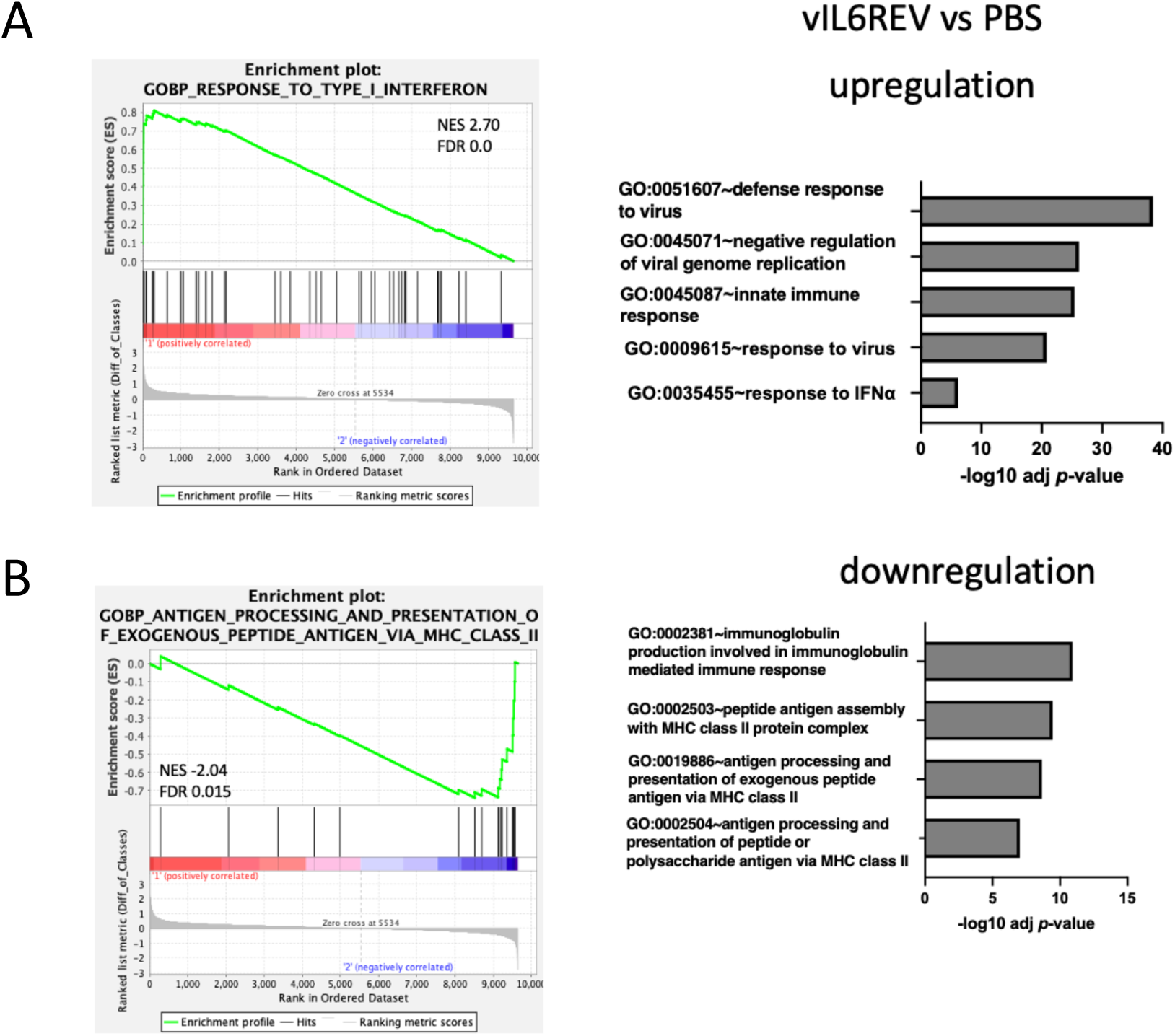
Gene Set Enrichment Analysis (GSEA) for vIL6REV-infected vs mock-infected macrophage transcriptomic landscape. The functional enrichment analysis of upregulated genes (A) and downregulated genes (B) in vIL6REV-compared to mock-infected (PBS treated) monocytes. DAVID functional enrichment analysis was performed, and the analysis results of top 5 enriched pathway are shown (Right). The representative results of GSEA are shown (Left).

